# SorCS2 deletion leads to altered neuronal lysosome activity

**DOI:** 10.1101/2021.04.08.439000

**Authors:** Sérgio Almeida, André M. Miranda, Andrea E. Tóth, Morten S. Nielsen, Tiago Gil Oliveira

**Affiliations:** Department of Biomedicine, Aarhus University, Aarhus, Denmark; Life and Health Sciences Research Institute (ICVS), School of Medicine, University of Minho, Braga, Portugal

## Abstract

Vps10p domain receptors are important for regulating intracellular protein sorting within the central nervous system and as such constitute risk factors for different brain pathologies. Here, we show that removal of SorCS2 leads to altered lysosomal activity in mouse primary neurons. SorCS2^-/-^ neurons show elevated lysosomal markers such as LAMP1 and acidic hydrolases including cathepsin B and D. Despite increased levels, SorCS2^-/-^ neurons fail to degrade cathepsin specific substrates in a live context. SorCS2-deficient mice present an increase in lysolipids, which may contribute to membrane permeabilization and increased susceptibility to lysosomal stress. Our findings highlight SorCS2 as an important factor for a balanced neuronal lysosome milieu.

## Introduction

Proper sorting of proteins to specific compartments is crucial for regular cell function. Therefore, abnormal protein sorting has been pointed out as the underlying cause of many brain pathologies like Alzheimer’s and Parkinson’s disease (Wang, Chan et al. 2013). The vps10p domain (Vps10p-D) receptors are a group of five structurally related type 1 transmembrane proteins that includes sortilin, SorLA, SorCS1, SorCS2 and SorCS3. As a common feature, the family members act as sorting receptors for different proteins within the central (CNS) and peripheral nervous system (PNS) (Willnow, Petersen et al. 2008). Predictably, the receptors have been shown to constitute risk factors for different neurodegenerative diseases such as frontotemporal dementia (FTD) in the case of sortilin (Philtjens, Van Mossevelde et al. 2018) and Alzheimer’s disease (AD) in the case of SorLA (Andersen, Reiche et al. 2005, Rogaeva, Meng et al. 2007). Sortilin, the archetype Vps10p-D receptor, has been pointed out as an endocytic and intracellular sorting receptor for the endo-lysosomal pathway in which it is responsible for the trafficking of different lysosomal proteins, such as acid sphingomyelinase (Ni and Morales 2006), cathepsins D and H (Canuel, Korkidakis et al. 2008) and progranulin (Hu, Padukkavidana et al. 2010). This lysosomal directed sorting is conveyed by the same sorting mechanism as observed with the mannose-6 phosphate receptors (M6PR), which are also responsible for delivery of newly synthesized hydrolases from the trans-Golgi network (TGN) to the late endocytic compartments (Nielsen, Madsen et al. 2001, Gary-Bobo, Nirdé et al. 2007, Mari, Bujny et al. 2008).

SorCS2 is a receptor of the same family, and belongs to the SorCS subfamily of receptors. It differs from sortilin by having a leucine rich domain and a Polycystic Kidney Disease (PKD) domain on the extracellular part (Hermey, Riedel et al. 1999) and is predominantly expressed in the CNS neurons, located mostly intracellularly, but it can also be found at the cell surface (Glerup, Olsen et al. 2014). SorCS2 has been associated with bipolar disorder (Baum, Akula et al. 2008, Ollila, Soronen et al. 2009), schizophrenia (Christoforou, McGhee et al. 2011) and attention deficit-hyperactivity disorder (ADHD) (Alemany, Ribasés et al. 2015) and different ligands have been identified such as neurotrophins for instance (Deinhardt, Kim et al. 2011, Anastasia, Deinhardt et al. 2013, Glerup, Olsen et al. 2014). Recently, SorCS2 has been implicated in different processes of the endocytic recycling pathway. By binding to cysteine transporter, excitatory amino acid transporter 3 (EAAT3), SorCS2 facilitates EEAT3 cell surface expression and enables neuronal oxidative stress defense (Malik, Szydlowska et al. 2019). Loss of SorCS2 leads to reduced EAAT3 cell surface expression due to a shift of the receptor from the recycling endosome to the late endosome bound for degradation. Additionally, SorCS2 was found to interact with the key component of the retromer complex Vps35, promoting the trafficking of NR2A subunit to the dendritic surface of medium spiny neurons (MSNs), showing that impaired SorCS2 activity contributes to motor coordination deficits in Huntington’s disease (HD) (Ma, Yang et al. 2017). Despite these recent advances, the effect of SorCS2 deficiency in late compartments of the endocytic pathway remains unknown.

Here, we show that the absence of SorCS2 leads to inversed modulation of early and late endocytic compartments in primary mouse neurons and mouse brain cortex. Additionally, we show that SorCS2 genetic ablation in mouse primary neurons impairs lysosomal degradation of cathepsin B substrates and renders the neurons more susceptible to lysosomal related stressors such as chloroquine (CQ) and alpha-synuclein (α-syn). Finally, SorCS2-deficient mice display an increase in lysolipids that may account for altered lysosomal milieu and subsequent decreased lysosomal degradation. Taken together, these findings describe a novel role for SorCS2 as a regulator of lysosomal environment and function.

## Results

### SorCS2 ablation leads to altered endo-lysosomal associated proteins in the mouse brain and primary neuronal cultures

We started by evaluating the effects of SorCS2’s absence on the different endo-lysosomal compartments in primary mouse neurons, such as Rab5 and EEA1 for early endosomes, Rab7 for late endosomes/multivesicular bodies and LAMP1 for lysosomes (**Fig. 1A**). Immunoblot analysis revealed higher levels of the early endosome marker Rab5, while the subsequent downstream compartment markers, EEA1 and Rab7, showed decreased levels in SorCS2^-/-^ neurons. Interestingly, the levels of the lysosome associated marker LAMP1 were also found to be increased in SorCS2-deficient neurons. Immunocytochemistry of hippocampal primary WT and SorCS2^-/-^ neurons analyzed by high content screening microscopy and scanR showed elevated numbers of LAMP1 positive vesicles in each SorCS2 ^-/-^ neuron and decreased number of EEA1 positive vesicles. This is in accordance with the immunoblot results and further shows a greater abundance of lysosomes and decreased intermediate compartments in the absence of SorCS2 (**Fig. 1B**).

**Figure 1.**
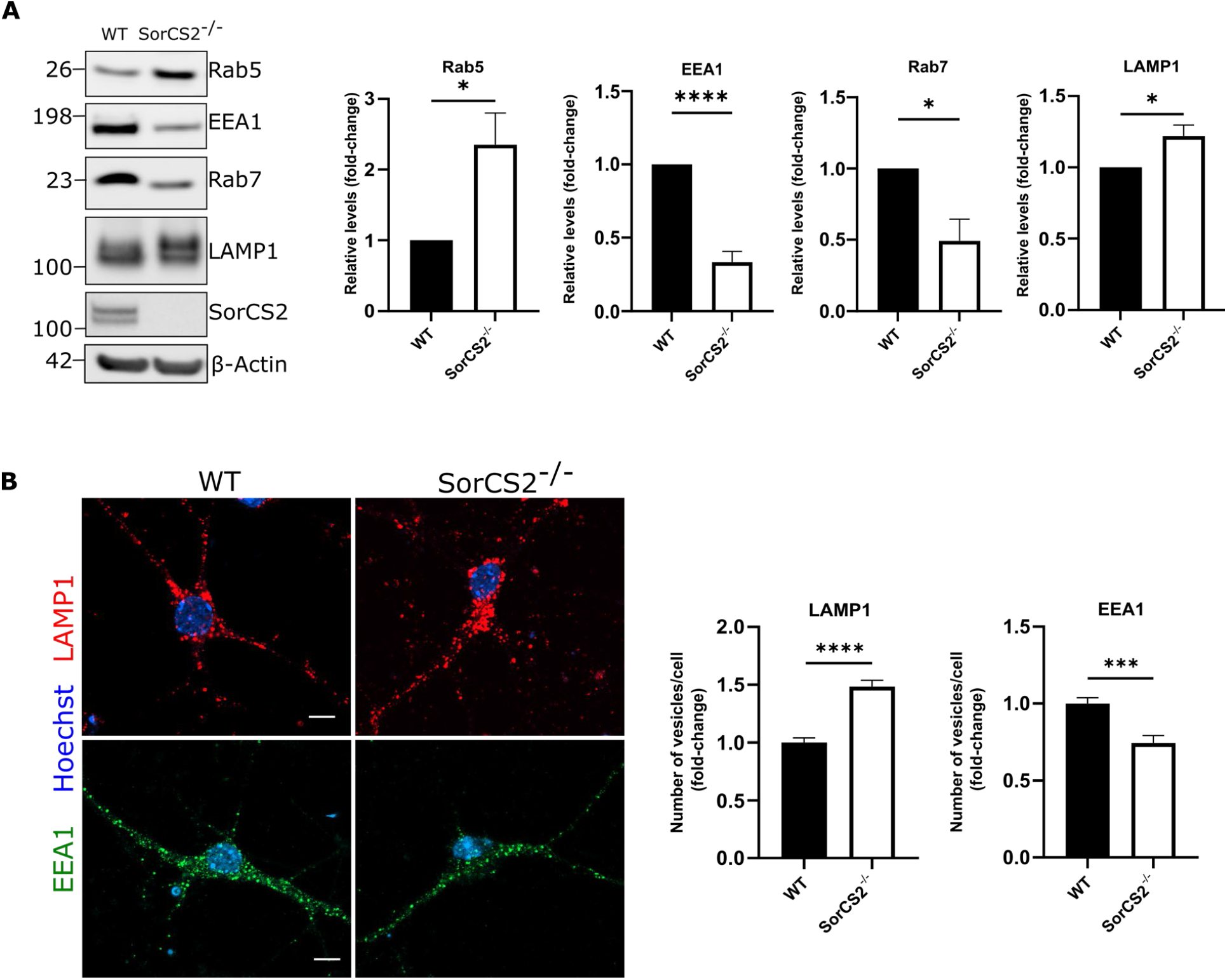
Absence of SorCS2 causes changes in different endo-lysosomal compartments. **A**. Western blot analysis of endo-lysosomal pathway associated proteins from WT and SorCS2^-/-^ primary neurons. β-actin served as loading control. Bar graphs denote average protein levels normalized to WT (mean ± SEM). *N*=6 independent neuronal preparations per genotype. *p<0.05, **p<0.01 (unpaired t test). **B**. Representative confocal images of LAMP1 (red) and EEA1 (green) immunostaining in WT (left) and SorCS2^-/-^ (right) mouse hippocampal neurons (single confocal z planes). Cells are counterstained with Hoechst. Scale bar, 10 µm. Bar graphs show scanR analysis of average LAMP1 and EEA1 vesicle numbers per neuron normalized to WT (mean ± SEM). *N*=454 and *N*=280 cells for LAMP1 staining of WT and SorCS2^-/-^ respectively and *N*=216 and *N*=160 cells for EEA1 staining of WT and SorCS2^-/-^ respectively, of three independent experiments. ****p<0.0001 (unpaired t test).

Next, we investigated the impact of SorCS2’s absence on the lysosomal hydrolases cathepsin B (CatB) and cathepsin D (CatD). CatD (and CatB) is produced in the biosynthetic pathway as precursor proCatD (46 kDA) and sorted to the endo-lysosomal compartment, where it is processed into a mature form (30 kDa) at acidic pH (Braulke and Bonifacino 2009). We found elevated levels of CatB and D mature forms, which correlated with increased number of lysosomes in neurons. Notably, we were able to assess a higher maturation of CatD based on the CatD/proCatD (**Fig. 2A**). This cathepsin maturation seemingly points to proper lysosomal function.

**Figure 2.**
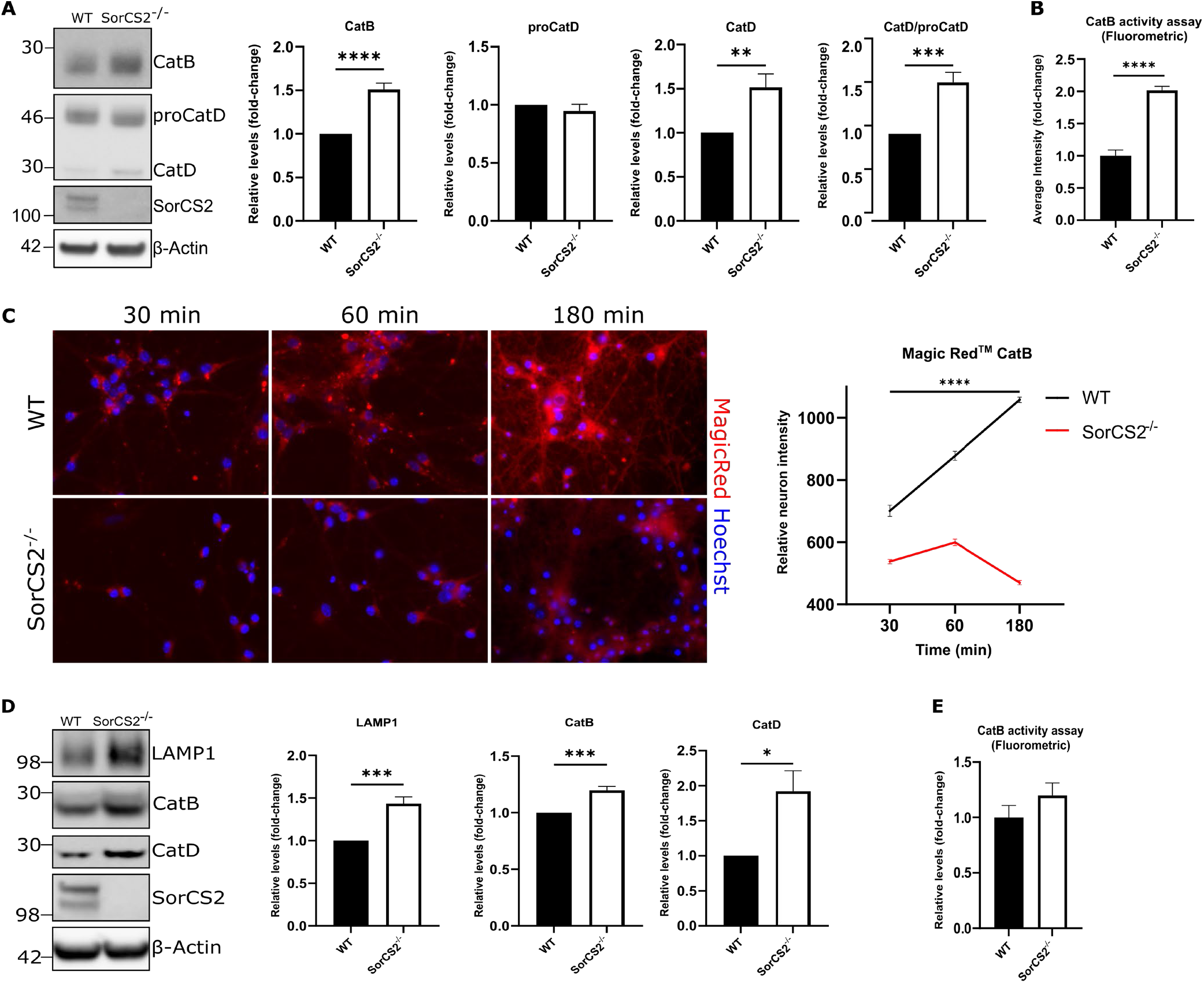
Cathepsin maturation and activity is altered in mouse SorCS2^-/-^ neurons and cortex. **A**. Western blot analysis of lysosome related proteins LAMP1 and mature Cathepsin B and D from WT and SorCS2^-/-^ primary neurons. β-actin served as loading control. Bar graphs denote average protein levels normalized to WT (mean ± SEM). *N*=6 independent neuronal preparations per genotype. *p<0.05, **p<0.01 (unpaired t test). **B**. Bar graph shows average fluorescent intensity resulting from abcam’s Cathepsin B activity assay kit (fluorometric) from WT and SorCS2^-/-^ primary cortical neurons, normalized to WT (mean ± SEM). *N*=4 independent neuronal preparations per genotype. ****p<0.0001 (unpaired t test). **C**. Live high content screening microscopy representative pictures of WT (top) and SorCS2^-/-^ (bottom) primary hippocampal neurons incubated with Magic Red™ Cathepsin B substrate for the designated time points (30, 60 and 180 min). Cells are counterstained with Hoechst. Graph denotes average neuron fluorescent intensity for both genotypes at specified time points (mean ± SEM). *N*=500 and *N*=613 cells at 30 min, *N*=569 and *N*=515 cells at 60 min, *N*=242 and *N*=502 cells at 180 min for WT and SorCS2^-/-^ respectively of three independent experiments. ****p<0.0001 (two-way ANOVA with Tukey’s multiple-comparisons test). **D**. Western blot analysis of lysosome related proteins LAMP1 and mature Cathepsin B and D from WT and SorCS2^-/-^ adult mouse cortex. β-actin served as loading control. Bar graphs denotes average protein levels normalized to WT (mean ± SEM). *N*=5 mice per genotype. *p<0.05, **p<0.01 (unpaired t test). **E**. Bar graph shows average fluorescent intensity resulting from abcam’s Cathepsin B activity assay kit (fluorometric) from 1-year-old WT and SorCS2^-/-^ mouse cortex, normalized to WT (mean ± SEM). *N*=10 and 9, WT and SorCS2^-/-^ mice respectively.

With the purpose of assessing the lysosomal degradative capacity, WT and SorCS2^-/-^ neurons were processed to measure CatB activity *ex vivo* (**Fig. 2B**). As expected, the increased mature CatB present in SorCS2^-/-^ neurons displayed increased signal when compared to WT neurons, as more enzymes will be able to degrade the substrate faster and produce higher signal. In addition, to evaluate CatB activity *in vitro*, we performed live imaging of primary neurons treated with Magic Red™ Cathepsin B, a membrane-permeable anionic compound that accumulates in acidic compartments and emits fluorescence upon cleavage by CatB. High content screening microscopy and scanR analysis at three different time points (30, 60 and 180 min) revealed time-dependent increase in fluorescence in WT neurons (**Fig. 2C**), suggesting increased load in late endocytic compartments. SorCS2^-/-^ neurons failed to produce the fluorescent product. As such, our results suggest that, despite elevated lysosomal number and increased cathepsin levels, CatB shows decreased processing *in vivo*, indicating possible changes in the lysosomal milieu.

Immunoblot analysis of cortex tissue from WT and SorCS2^-/-^ adult mice further confirmed the observations in neurons regarding increased levels in lysosomal associated proteins (LAMP1, CatB and CatD mature forms) (**Fig. 2D**). Additionally, we performed ex vivo CatB activity assay in brain hippocampus from 1-year-old mice WT and SorCS2^-/-^. SorCS2 deletion in mice resulted in elevated CatB activity when compared to WT, although not statistically significant (**Fig. 2E**).

### SorCS2^-/-^ neurons are more susceptible to lysosomal stress

Since there is an increase in lysosomes upon SorCS2 ablation, we evaluated the neuronal response to lysosomal stressors, such as CQ and α-syn. Cultured WT and SorCS2^-/-^ primary mouse neurons were incubated with 50 µM CQ, known for inhibiting lysosomal acidification and impair autophagy (Seglen, Grinde et al. 1979, Poole and Ohkuma 1981, Mizushima, Yoshimori et al. 2010) (**Fig. 3A**). SorCS2^-/-^ neurons show higher levels of cleaved caspase-3 when compared to WT neurons, indicating a higher rate of apoptosis. Autophagy marker LC3 was probed as a control for lysosomal neutralization and indeed, LC3-II accumulates to the same extent upon treatment with CQ in both genotypes, suggesting that apoptosis was independent of autophagy machinery. Higher susceptibility was also confirmed using LIVE/DEAD™ Cytotoxicity assay in which SorCS2^-/-^ neurons show a higher percentage of dead cells when submitted to lysosomal stress (**Fig. 3B**).

**Figure 3.**
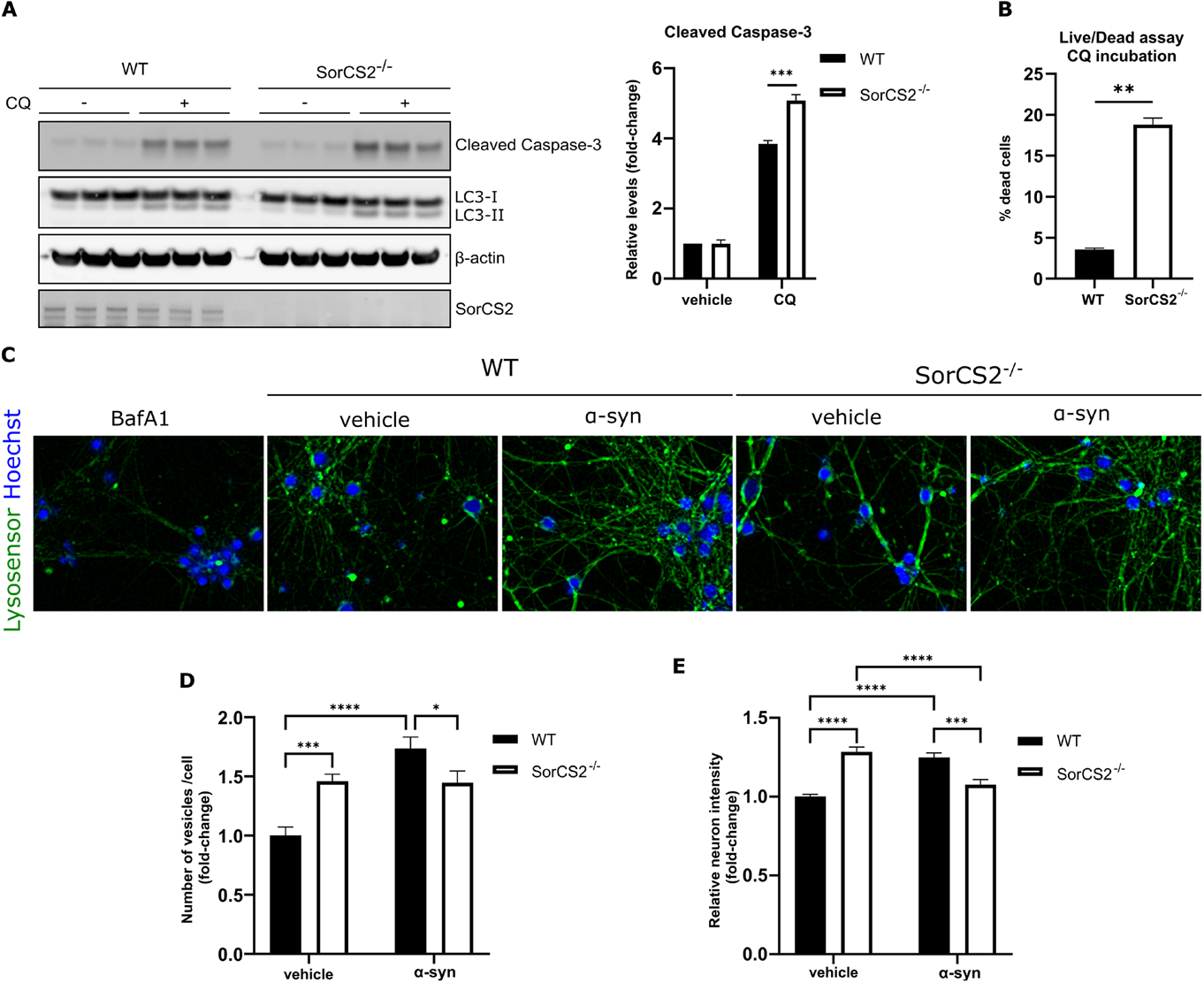
SorCS2^-/-^ primary neurons are susceptible to lysosomal stress. **A**. Western blot analysis of cleaved caspase-3 levels in WT and SorCS2^-/-^ primary cortical neurons treated with vehicle or CQ at 50 µM for 24h. LC3 serves as lysosomal impairment control and β-actin as loading control. Bar graphs denotes average protein levels normalized to WT (mean ± SEM). *N*=3 independent neuronal cultures. ***p<0.001 (two-way ANOVA with Tukey’s multiple-comparisons test). **B**. Bar graph shows percentage of dead WT and SorCS2^-/-^ primary cortical neurons treated with vehicle or CQ at 50 µM for 24h (mean ± SEM). *N*=2 independent neuronal cultures. **p<0.01 (unpaired t test). **C**. Live high content screening microscopy representative pictures of WT and SorCS2^-/-^ primary hippocampal neurons incubated with Lysosensor™ Green DND-189 and pre-incubated with either BafA1 for at 100 nM for 45 min, vehicle and monomeric α-syn for 24 h at 14 µg/ml. Cells are counterstained with Hoechst. **D and E**. scanR analysis from images in **C. D**. Bar graph denotes average number of vesicles per neuron for both genotypes when treated with either vehicle or αα-syn, normalized to the WT (mean ± SEM). *N*=150 and *N*=289 cells for vehicle treated and *N*=239 and *N*=133 cells for α-syn treated WT and SorCS2^-/-^ respectively of three independent experiments. *p<0.05, ***p<0.001, ****p<0.0001 (two-way ANOVA with Tukey’s multiple-comparisons test). **E**. Graph denotes average vesicle fluorescent intensity for both genotypes when treated with either vehicle or α-syn, normalized to the WT (mean ± SEM). *N*=150 and *N*=289 cells for vehicle treated and *N*=239 and *N*=133 cells for α-syn treated WT and SorCS2^-/-^ respectively of three independent experiments. ****p<0.0001 (two-way ANOVA with Tukey’s multiple-comparisons test).

It has been documented that lysosome biogenesis is elevated when cells are exposed to exogenous α-syn, as this protein is mainly cleared by the endo-lysosomal pathway (Hoffmann, Minakaki et al. 2019). Therefore, to evaluate the lysosomal response to robust levels of α-syn exposure in the absence of SorCS2, neurons were incubated with 14 µg/ml of monomeric α-syn for 24 h, and then stained with Lysosensor™ Green DND-189, a pH sensitive probe (**Fig. 3C**). Neurons were subsequently analyzed using high content screening microscopy. We confirmed an increase in lysosomal numbers, as expressed by the number of punctate structures (**Fig. 3D**), but also higher acidification of vesicles when SorCS2 is ablated, as expressed by increased fluorescence intensity (**Fig. 3E**). However, upon α-syn treatment, while lysosomal numbers increased in WT neurons, KO neurons were unable to elevate their lysosomal numbers and showed decreased vesicle intensity compared to baseline. This loss in acidification might be due to lysosomal membrane permeabilization (LMP) and consequent leakage (Jiang, Gan et al. 2017).

### Removal of SorCS2 increases lysolipid production in mouse cortex

To gain further insight into the potential effect of SorCS2 deletion in the endo-lysosomal compartments, we performed a mass spectrometry lipidomic analysis in the brain cortex of adult WT and SorCS2^-/-^ mice. We identified 19 lipid classes that were further divided into 147 lipid species (**Fig. 4A**). Of all the lipid classes, lysolipids lysophosphatidic acid (LPA) and lysophosphatidylserine (LPS) were the most significantly elevated. While these are of very reduced abundance (0.02-0.05% of all lipids measured), lysolipids can be synthesized in late endocytic compartments and, given their physical structure, they can increase the instability of phospholipid membranes and consequently increase their permeabilization (Harayama and Riezman 2018). Lysolipids may act as intermediates for glycerophospholipid synthesis, but may also be produced by action of phospholipases, namely PLA, of which there are many isoforms differently localized (D’Arrigo and Servi 2010).

**Figure 4.**
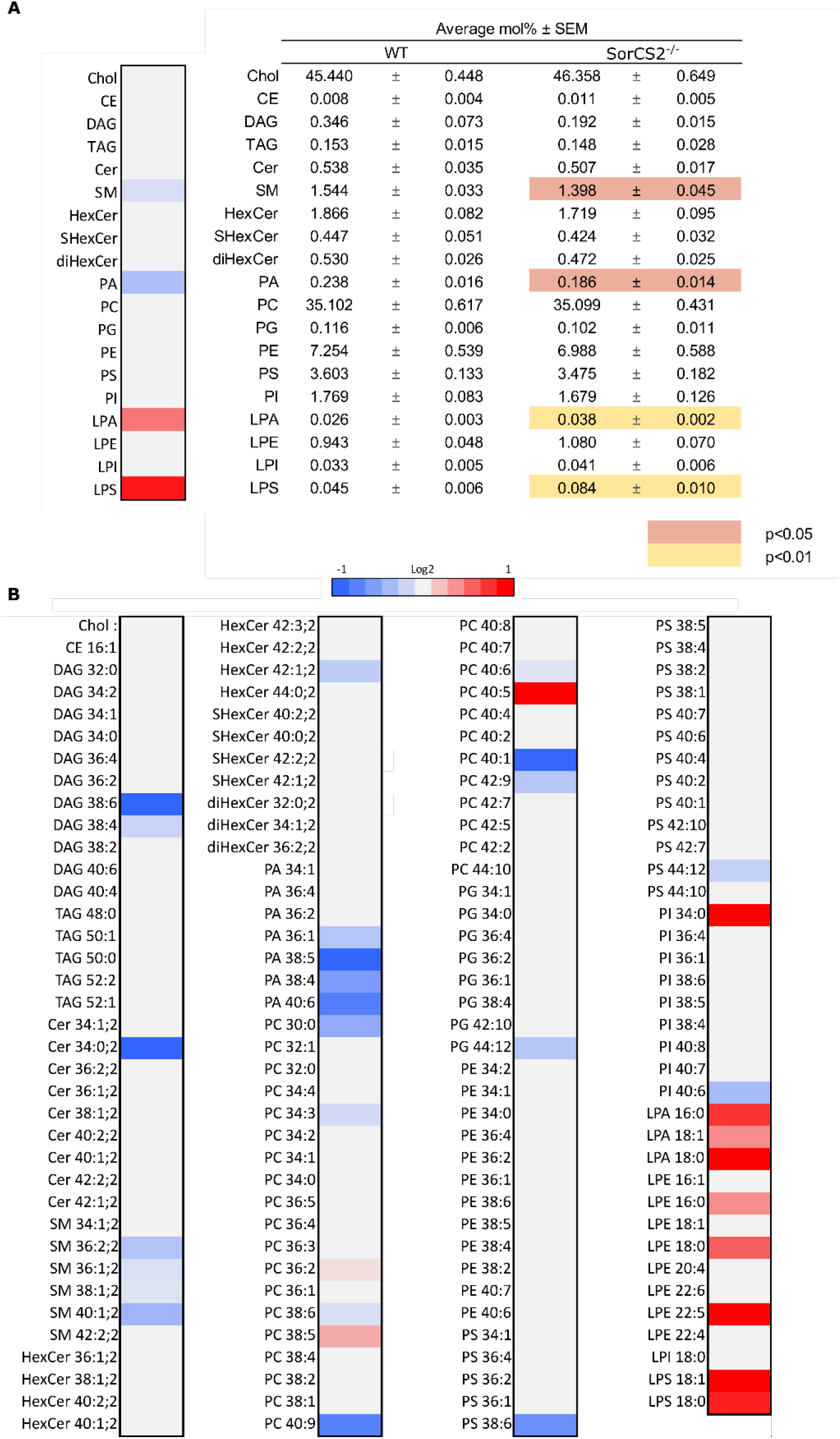
Lipidomics analysis of WT and SorCS2^-/-^ mouse cortex. MS/MS analysis of **A**. Lipid classes and **B**. Lipid species displayed as a Heatmap from 1-year-old WT and SorCS2^-/-^ mouse cortex. For lipid nomenclature, consult Methods section. Values are expressed as average mol% of total lipid measured, normalized to WT (mean ± SEM, *N*=10 and 9, WT and SorCS2^-/-^ mice respectively)

A significant reduction is also observed in phosphatidic acid (PA) and sphingomyelin (SM). To further amplify the study, individual lipid species were investigated (**Fig. 4B**). A significant impairment in 35 different lipid species was seen. Curiously, we find decreased sphingomyelin and ceramide species, while no changes were detected in more complex sphingolipids, suggesting decreased synthesis or increased degradation. Interestingly, the precursor for these species, sphingosine has been reported to induce LMP in increased concentrations (Kågedal, Zhao et al. 2001). Additionally, changes are observed in multiple phosphatidylcholine (PC) and phosphatidic acid (PA) species.

## Discussion

Lysosomes are central organelles crucial for proper cell function and lysosomal dysfunction has been implicated in many neurodegenerative diseases (Zhang, Sheng et al. 2009). Defects in trafficking of different proteins may result in importer sorting of different components which will cause changes in the different compartments of the endocytic pathway and ultimately impair the lysosome degradative capacity (Ghosh, Dahms et al. 2003). SorCS2, being a transmembrane sorting receptor, has been reported to act as a co-receptor and together with TrkB facilitate BDNF signaling and TrkB translocation to the postsynaptic densities (Glerup, Bolcho et al. 2016). Together with the recent discoveries of SorCS2’s involvement in the endocytic recycling pathway and therefore controlling cell surface exposure of receptors (Ma, Yang et al. 2017, Malik, Szydlowska et al. 2019, Yang, Ma et al. 2021), it is expected that its absence will affect the different endocytic compartments.

Here, we show that the absence of SorCS2 leads to an increase in lysosomes and, even though cathepsins are shown to mature, their activity in live neurons is seemingly impaired, pointing towards changes in the lysosomal milieu such as altered pH, altered trafficking of co-factors or accumulation of enzymatic inhibitors, which may be unfavorable to protease action. Indeed, despite increased lysosomal acidification, SorCS2-deficient neurons show an abnormal reaction when faced with lysosomal stressors such as CQ and α-syn. SorCS2^-/-^ neurons show a higher death rate in response to CQ, are unable to raise their lysosomes in response to exogenous α-syn and present decreased lysosomal acidification. This may suggest that elevated levels of LAMP1 and lysosomal acidification at steady-state levels already indicate an underlying defect and upregulated response to attempt to establish homeostasis, which is unelastic in the context of additional stress. Importantly, exogenous exposure to α-syn has been shown to induce lysosomal rupture following endocytosis in neuronal cell lines and induce a CatB dependent increase in reactive oxygen species (ROS) (Freeman, Cedillos et al. 2013, Jiang, Gan et al. 2017).

Lipidomic analysis report specific changes in lipid classes and species, suggesting a relative localized effect caused by the absence of SorCS2. The increase observed in lysolipids will affect phospholipid membranes by altering their shape and constitution, further supporting the vulnerability of the late endocytic compartments (Fuller and Rand 2001, Chevallier, Chamoun et al. 2008, Bissig and Gruenberg 2013). This might be mediated by phospholipases such as PLA2G4A/cPLA2, which has been shown to promote LMP and inhibition of autophagy flux leading to neuronal dysfunction and eventually cell death in mice with traumatic brain injury (Sarkar, Jones et al. 2020). In order to investigate the possible LMP, future studies should be conducted in WT and SorCS2^-/-^ neurons to directly image effectors of lysosomal membrane repair, such as early recruitment of galectin and subsequent autophagy machinery upon α-syn incubation (Aits, Kricker et al. 2015). It will be interesting to address how lysosomal function is affected, whether there is decreased formation of engulfing autophagosomes or decreased processing of these structures. Notably, when incubated with CQ, SorCS2^-/-^ neurons reveal the same extent of LC3-II accumulation as control neurons. This suggest others aspects than autophagy are impaired, hence the impaired levels of specific endocytic modulators (e.g. Rab7 and Rab5).

SorC2 deficient mice display behavioral changes such as impaired formation of long-term memory, increased risk taking, ADHD-like behavior (Glerup, Olsen et al. 2014, Glerup, Bolcho et al. 2016, Olsen, Wellner et al. 2021) and social memory deficit (Yang, Ma et al. 2021). Curiously, some of these phenotypes have been reported in phospholipase D1 (PLD1) deficient mice, which also show decreased levels of the PA lipid (Santa-Marinha, Castanho et al. 2020) as our lipidomics results on SorCS2^-/-^. PLD1 is a major source of PA and it is primarily associated with the endosomal system where it was suggested to be recruited by class III PI 3-kinase Vps34 (Dall’Armi, Hurtado-Lorenzo et al. 2010), which is a major regulator of endolysosomal and autophagic functions (Backer 2016). Finally, disruption of neuronal Vps34 function impairs autophagy, lysosomal degradation and lipid metabolism, affecting different cellular compartments and causing endolysosomal membrane damage (Miranda, Lasiecka et al. 2018) similar to what we observe in SorCS2^-/-^ neurons.

SorCS2 mediates recycling of different receptors to the cell surface, documented to rely on the binding to Vps35 (Ma, Yang et al. 2017). The component of the retromer complex Vps35 is a major regulator of endosomal sorting (Burd and Cullen 2014) and several studies have shown that reduced levels of Vps35 affect the traffic of cation-independent (CI)-M6PR (Arighi, Hartnell et al. 2004, Seaman 2004, Seaman 2007, Wassmer, Attar et al. 2007, Bulankina, Deggerich et al. 2009, Miura, Hasegawa et al. 2014, Hirst, Itzhak et al. 2018, Seaman 2018). Furthermore, loss of retromer Vps35 subunit impairs targeting and processing of M6PR-depedent hydrolases, causing ultrastructural alterations and compromised lysosomal function, leading to impaired autophagy (Cui, Carosi et al. 2019). Interestingly, phospholipase *iPLA2-VIA* (the fly homolog of PLA2G6) has been shown to bind to Vps35 and enhance retromer function to promote protein and lipid recycling, where loss of this phospholipase impairs retromer and lysosomal function in flies (Lin, Lee et al. 2018). Future experiments should address the levels and localization of Vps35 and PLA2G6 under the absence of SorCS2.

In summary, we conclude that SorCS2 may act as an indirect mediator of neuronal lysosomal function and a missing piece in the regulation and correct sorting of the retromer complex subunit Vps35. In the absence of SorCS2, Vps35 is mistrafficked leading to abnormal phospholipase activity and further irregular sorting of components to the lysosomes, compromising equilibrium. The consequent decrease in acidic hydrolase activity altered the lysosomal milieu causing higher vulnerability to lysosomal stressors. The data collectively suggests promising avenues for the investigation of the neuronal endocytic pathway mechanisms regulated by SorCS2.

## Materials and Methods

### Reagents and antibodies

Bafilomycin A1 (100 nM, Merck, B1793), CQ diphosphate (50 µM, Merck, C6628), mouse monomeric α-synuclein (kind gift from Poul Henning Jensen, Aarhus University). Antibodies: α-synuclein (BD Biosciences, 6107887, 1:1000 in WB), β-actin (Merck, A5441, 1:5000 in WB), Cathepsin B (Cell Signaling, D1C7Y, 1:1000 in WB), Cathepsin D (Abcam, ab75852, 1:1000 in WB), Cleaved Caspase-3 (Cell Signaling, 9661, 1:1000 in WB), EEA1 (Cell Signaling, C45B10, 1:1000 in WB and 1:100 in ICC) EGFR (EMD Millipore, 06-847, 1:1000 in WB and 1:100 in ICC), LAMP1 (Abcam, ab24170, 1:1000 in WB and 1:100 in ICC), LC3 (Novus Biologicals, NB600-1384, 1:1000 in WB), Rab5 (Synaptic Systems, 108011, 1:1000 in WB), Rab7 (Abcam, ab137029) and SorCS2 (R&D systems, AF4237, 1:1000 in WB and 1:100 in ICC).

### Animals

1 year-old female C57BL/6j and *SorCS2*^*-/-*^ mice were used for experiments. SorCS2^-/-^ on a C57BL/6j background were described before (Glerup, Olsen et al. 2014). The mice were bred and housed at the Animal Facility at Aarhus University. The mice were housed in groups and kept under pathogen-free conditions with a 12 h light/ 12 h dark schedule with food and water *ad libitum*. All procedures involving mice were approved by the Danish Animal Experiment Inspectorate (license 2017-15-0201-01203) and complied with Danish and European regulations (directive 2010/63/EU) concerning experimentation and care of experimental animals.

### Cell culture

Primary cortical and hippocampal neurons were obtained from wild-type and SorCS2^-/-^ postnatal day 0 mouse pups of either sex (Glerup, Olsen et al. 2014). Pups were euthanized by decapitation, brains removed, and hippocampi or cortex dissected into ice-cold Leibowitz’s L-15 medium (Life technologies, 11415049). Tissue was dissociated for 30-40 min using 20 U/mL pre-activated papain (Bionordica, WBT-LS003126) and washed in DMEM containing 0.01 mg/mL DNase I (Merck, DN25) and 10% FBS before being triturated in the same buffer. Following trituration, DMEM was substituted with Neurobasal-A Medium (Gibco, 10888022) supplemented with 2% B-27 (Gibco, 17504044), 2 mM GlutaMAX (Gibco, 35050), 100 µg/mL primocin (Invivogen, ant-pm-2), 20 µM floxuridine, and 20 µM uridine (Merck, F0503, U3750). Cortical neurons were seeded onto pre-coated wells with poly-L-lysine (Merck, 15140) and laminin (invivogen, 23017-015) and hippocampal neurons were seed into µ-Slide 8 well (ibidi, 80826) pre-coated with poly-D-lysine (Merck, P7886). Neurons were incubated at 37°C for 7 days prior to assaying with medium change every second day.

### Survival assays

WT and SorCS2^-/-^ primary cortical neurons were plated into 12-well plates and 96-well black plate with clear bottom (Corning Incorporated, Costar, 3603). On day in vitro 7 (DIV7), neurons were incubated with 50 µM CQ diphosphate (Merck, C6628) dissolved in water for 24 h. On the next day, neurons were harvested for immunoblotting or incubated with calcein-AM and ethidium homodimer-1 (LIVE/DEAD™ Viability/Cytotoxicity Kit, Thermofisher, L3224), following the manufacturer’s instructions. Fluorescence intensity was read using CLARIOstar plus plate reader (BMG labtech) with appropriate filters.

### Immunofluorescence

Cultured primary hippocampal neurons plated onto µ-Slide 8 well (ibidi, 80826) were fixed in 4% paraformaldehyde, 4% sucrose (Merck, S0389) in D-PBS for 15 min and permeabilized with 0.05% saponin (Merck, 47036) in D-PBS supplemented with 1% bovine serum albumin (BSA, Merck, 05470). Primary antibodies diluted in blocking buffer were incubated with the cells at RT for 1 h, followed by a three times 5 min washing step with blocking buffer. Alexa-conjugated secondary antibodies were sequentially incubated for 1 h in the same buffer, protected from light. For mounting, ibidi Mounting Medium with DAPI (ibidi, 50011) were added to the cells in the µ-Slide 8 well.

### Confocal Microscopy

The immunostained samples were analyzed on a Zeiss confocal LSM 710 microscope using 63x/1.2 W Korr (water immersion correction ring) objectives. Appropriate filters were used upon excitation of the different fluorophores to match their maximum fluorescence emission. Processing of the acquired images was performed in ImageJ. All images compared were subjected to similar brightness and contrast adjustments.

### Live cell microscopy

For the live cell assays, WT and SorCS2^-/-^ primary hippocampal neurons were plated in µ-Slide 8 well and incubated as described before. On DIV6, neurons were incubated with either vehicle or 14 µg/ml homemade monomeric α-synuclein for 24 h. On DIV7, neurons were incubated with prewarmed media containing 1 μM LysoSensor Green DND-189 (Thermofisher, L7535) and 0.125 μg/ml Hoechst 33342 (Thermofisher, H3570) for 15 min at 37 °C. Bafilomycin A1 (BafA1) was used prior to LysoSensor Green DND-189 incubation at 100 nM for 45 min as a control for acidification. Cells were washed twice with D-PBS and kept in FluoroBrite DMEM media (Gibco, A1896701) supplemented with 1% GlutaMAX for imaging. Method described before (Toth, Nielsen et al. 2019).

For the Cathepsin B live activity assay, Magic Red® Cathepsin B substrate (ImmunoChemistry Technologies, 937) was added to the neurons following manufacturer’s instructions and incubated for 30, 60 and 180 min prior to imaging. At the end of the time points neurons were washed twice in D-PBS and incubated with FluoroBrite DMEM media supplemented with 1% GlutaMAX and 0.125 μg/ml Hoechst 33342 (provided with the Magic Red® Cathepsin B kit). Cells were then imaged in a microscope equipped with a live cell imaging chamber in humidified atmosphere with 5% CO2 at 37 °C. Analysis of the different parameters was performed with a high content screening fluorescence microscope (Olympus BX73 microscope) fitted with the correct filter set as described below.

### High content screening analysis

Following immunocytochemistry, LysoSensor dye and Magic Red Cathepsin B activity substrate uptake, images for high content screening were obtained with the Olympus automated Scan^R high content imaging station based on an Olympus BX73 microscope, with a 40×/0.9 NA air objective or 60x/0.9 NA air objective, triple-band emission filter for DAPI, FITC and Cy3, and a Hamamatsu camera (C8484-05G). Image analysis was performed using scanR image and data analysis software for Life Science (Münster, Germany). Briefly, single-layer images were background-corrected and edge-detection algorithm was applied to segment subcellular structures based on detection of gradient intensities of the chosen color channel. The software segmented subcellular structures independently if a closed connecting line (edge) could be drawn around them and their area was larger than 0.05 μm2 independently of their shape. Images with artefacts or out of focus were manually excluded. The total number of vesicles was normalized to the number of nuclei before making comparison among the adjacent groups. Number and mean fluorescent intensity of vesicles from 100 to 500 cells for each group were analyzed (Toth, Nielsen et al. 2019).

### Protein biochemistry and immunoblotting

Cells were lysed and scraped in homemade Radioimmunoprecipitation assay buffer (RIPA) ((50 mM Tris pH 7.4, 150 mM NaCl, 1 % Triton X-100 (Merck, T9284), 2 mM EDTA, 0.5 % Sodium-deoxycholate (Merck, D6750), 0.1 % sodium dodecyl sulfate (SDS) (Merck, cat. no. L6026)) supplemented with 1x cOmplete™ Protease Inhibitor Cocktail (Merck). Homogenates were centrifuged at top speed for 5 min at 4° C and supernatant was processed for protein dosage (BCA, Pierce). Samples diluted to equal concentration were prepared with NuPAGE LDS sample buffer and 0.02 M dithiothreitol (DTT), heated for 10 min at 95°C and loaded into NuPAGE 4-12% Bis-Tris gels; separation was carried out using MES running buffer (Thermofisher). Protein transfer was performed to nitrocellulose membranes using iBlot Transfer Stack (Thermo Fisher, IB301032 and IB301031) according to manufacturer’s protocol. Membranes were blocked for 1 h at room temperature with blocking buffer (0.01 M Tris-Base, 0.15 M NaCl, 0.05 % Tween-20, 5 % skimmed milk powder and 0.002 % azide, Ampliqon, AMPQ52300) and subsequently incubated with primary antibodies diluted in the same buffer overnight at 4°C. HRP-conjugated secondary antibodies were incubated for 1 h at room temperature and membrane development was performed using Enhanced Chemiluminescence (ECL) Western Blotting substrate (GE healthcare, RPN2106) or ECL Prime Western Blotting substrate (GE healthcare, RPN2236). Chemiluminescent signal was imaged with Fujifilm Las3000 imaging apparatus and analyzed using Fujifilm MultiGauge software.

### Cathepsin B activity assays kits (Fluorometric)

Cathepsin B activity was assessed in mouse primary neuronal samples and mouse brain hippocampus using Fluorometric Cathepsin B activity assay kit (abcam, 65300) following the manufacturer’s protocol. Fluorescence intensity was read using CLARIOstar plus plate reader (BMG labtech).

### Lipidomics

1 year-old WT and SorCS2^-/-^ mice were euthanized by cervical dislocation. Cortex was dissected and snap frozen in liquid nitrogen and stored at -80° C. The lipidomics analysis was performed by the Lipidomics Core Facility at the Danish Cancer Society Research Center. Tissue was mixed with chloroform:methanol 30 10:1 (v/v) and 10x internal lipid standards containing known concentrations of various lipids followed by shaking and centrifugation. The lower phase contains the nonpolar lipids (cholesterol (Chol), cholesteryl ester (CE), ceramide (Cer), HexoseCeramide (HexCer), sphingomyelin (SM), diacylglycerol (DAG), triacylglycerol (TAG), lysophosphatidylethanolamine (LPE), lysophosphatidylcholine (LPC), phosphatidylglycerol (PG), phosphatidylethanolamine (PE), phosphatidylcholine (PC). Chloroform:methanol 2:1 (v/v) was added to the upper phase to isolate polar lipids followed by shaking and centrifugation. The lower phase contains the polar lipids (diHexoseCeramide (diHexCer), lysophosphatidic acid (LPA), lysophosphatidylinositol (LPI), lysophosphatidylserine (LPS), phosphatidic acid (PA), phosphatidylinositol (PI), phosphatidylserine (PS). The lipids were dried in a vacuum centrifuge, dissolved in chloroform:methanol 1:2 (v/v) and transferred to a 96-well plate. The samples were analyzed using a robotic nanoelectrospray ion source coupled to the MS/MS on Q Exactive Hybrid Quadrupole-Orbitrap Mass spectrometer. Lipids were identified based on the unique mass-to-charge ratio (m/z) of the intact precursor ion and the subsequent

MS/MS spectra. Quantification was performed by comparing the peak intensity with the peak intensity of the internal standard lipid with a known quantity.

### Statistical analysis

Statistical analysis was performed using Prism software (Graphpad). All data are given as mean ± s.e.m. for a given *N* of biological replicates. Each mouse (or biological replicate) represents a statistically independent experimental unit, which was treated accordingly as an independent value in the statistical analysis. Comparison between two experimental groups was done through a two-tailed unpaired t-test. Two-way ANOVA followed by Tukey’s multiple comparisons test was performed for analysis of additional experimental groups and time-course experiments.

## Acknowledgements

We acknowledge the invaluable technical help provided by Mette Singers, Mathias Ollendorff and the AU Health Bioimaging core facility (Department of Biomedicine, Aarhus University). We are grateful to Prof. Poul Henning Jensen (Department of Biomedicine, Aarhus University) for kindly providing mouse α-syn protein and to Mesut Bilgin (Lipidomics Core Facility at the Danish Cancer Society Research Center) for Lipidomics sample processing. Sérgio Almeida is funded by a PhD fellowship from the PhD School at the Health Faculty, Aarhus University.

## Author contributions

Sérgio Almeida, André Miranda and Tiago Gil Oliveira developed the concept and designed the study. Sérgio Almeida and Andrea Tóth performed experiments. Sérgio Almeida and André Miranda analyzed the data. Sérgio Almeida drafted the manuscript. Morten Nielsen provided resources. Sérgio Almeida and André Miranda wrote jointly the final version of the manuscript. All authors critically revised the manuscript and approved the final version.

